# 3D Light Sheet Microscopy Reveals Aging Osteoarthritis-associated Joint-innervated Nerve Remodeling in Mouse Knee Joints

**DOI:** 10.64898/2026.01.09.698640

**Authors:** Paromita Kundu, Brian Wise, Emma Norris, Mark James, Yaxin Zhang, Jennifer Jonason, Mark Buckley, Whasil Lee

## Abstract

**Objectives:** Osteoarthritis (OA) is a prevalent age related joint disease-causing chronic pain. This study investigated how nociceptive and sympathetic nerve innervation in the mouse knee joint changes with age and OA progression, and how these changes relate to pain and disease severity.

**Methods:** Thirty-eight mice were assigned to four groups: young male (M_y_), aged male (M_a_), young female (F_y_), and aged female (F_a_). Pain sensitivity was evaluated via Pressure Application Measurement (PAM), and joint damage was graded using OARSI scoring. CGRP and PIEZO2 expressions in dorsal root ganglia (DRG) were also assessed. We employed iDISCO tissue clearing and 3D light sheet fluorescence microscopy to visualize total (PGP9.5⁺), nociceptive (CGRP⁺), and sympathetic (TH⁺) nerve fibers in anterior regions of mouse knee joints. A MATLAB-based tool quantified nerve architecture.

**Results:** The M_a_ group displayed the highest OA severity and markedly lower mechanical withdrawal thresholds compared with M_y_ (H = 11.59, ε² = 0.64), suggesting an age effect on pain-related behavior. This phenotype was accompanied by a higher total PGP9.5⁺ nerve fiber density in the knee joint (mean difference −0.3127, 95% CI −0.4551 to −0.1704) and increased CGRP⁺ nociceptive innervation. In contrast, female mice showed no age-dependent change in PAM withdrawal thresholds, consistent with preserved cartilage integrity and stable OARSI scores, and no detectable age-related differences in PGP9.5⁺ or CGRP⁺ innervation. The TH⁺ sympathetic fiber distribution was comparable across sexes and ages. Consistent with joint-level findings, DRG analyses demonstrated increased CGRP and PIEZO2 expression in the M_a_ group, whereas females exhibited no significant change.

**Conclusion:** Enhanced nociceptive but not sympathetic nerve remodeling in knee cavities is associated with increased OA severity and knee pain in the M_a_ group. These findings emphasize the role of peripheral sensory plasticity in OA pain and demonstrate the value of 3D imaging for visualizing neuroanatomical changes in joint disorders.

## Introduction

Osteoarthritis (OA) is a common degenerative joint disease that primarily affects the elderly, leading to chronic pain and functional limitations (1). Globally, around 240 million people suffer from symptomatic OA, with persistent pain as the most debilitating symptom (2, 3). For decades, OA has ranked among the leading causes of disability worldwide (4). The onset of OA-related pain is thought to be driven by a combination of cartilage degradation, subchondral bone remodeling, and inflammation of the synovium and periosteum (5). Notably, OA-associated joint pain exhibits marked sex differences. Accumulating evidence highlights a sexual dimorphism in nociception and pain processing, driven by hormonal regulation, genetic variation, and sex-specific behaviors (6, 7). The impact of aging and OA on spatially distinct knee joint innervation remains poorly defined, reflecting the lack of comprehensive joint-wide neural mapping. (8).

Earlier studies using light and electron microscopy in animal models, such as cryosections of cat knee joints, identified dense sensory and sympathetic innervation in the periosteum, menisci, ligaments, and subchondral bone (9). The composition of nerves in these tissues is dominated by sensory nerves, of which approximately 80% are nociceptors, highlighting their critical role in pain perception and joint function (10, 11). Research has also demonstrated that neuronal remodeling leads to the sprouting of both sensory and sympathetic axons in damaged tissue, contributing to pain (12, 13). Numerous studies have investigated nerve fiber distribution in knee joints using traditional histological and immunohistochemical staining of thin tissue sections to better understand pain mechanisms in OA (14). While staining thin, two-dimensional (2D) tissue cross-sections through the knee joint has provided keen insight, these studies fail to capture the three-dimensional (3D) spatial organization and connectivity of nerves within the complex structure of the knee joint. This limitation may hinder a comprehensive understanding of pain mechanisms in OA, as pain perception is influenced by the intricate network of nerves and their interactions with surrounding tissues.

To address this challenge, volume imaging technologies have emerged at the forefront of contemporary systems biology research. The immunolabeling-enabled three-dimensional imaging of solvent-cleared organs (iDISCO) technique is an advanced tissue-clearing method that facilitates high-resolution, 3D imaging of biological structures (15, 16). iDISCO has proven effective for tracing nerve fibers within mouse musculoskeletal tissues (17, 18). Recently, Ko et al. implemented a workflow of tissue clearing and light sheet microscopy to visualize knee-innervating nerve fibers in mouse joints (17). In this regard, labeling nerve-specific proteins allows to map the spatial distribution of nerves in the context of OA. However, the diverse composition of musculoskeletal tissues, including bone, articular cartilage, and muscle presents considerable challenges for clearing-enabled light sheet microscopy. To date, no study has evaluated how 3D nerve distribution changes across the entire knee joint in aging-associated OA.

We hypothesize that aging-associated OA involves pathological remodeling of knee joint innervation that contributes to joint pain. To evaluate this hypothesis, we used 3D light-sheet fluorescence microscopy with custom MATLAB-based spatial analysis to characterize sex specific nerve innervation patterns in the aging mouse knee, with emphasis on medial–lateral distribution. Analysis was performed within a standardized (anatomical region of interest) ROI spanning the full medial–lateral width of the joint and extending from the inferior patellar pole to the proximal tibial epiphysis. We further conducted a comprehensive set of experiments, including the pressure application measurement (PAM) to assess mechanical allodynia as a behavioral readout of pain sensitivity, OARSI scoring to evaluate cartilage degeneration, and immunohistochemistry to examine the expression of pain-associated markers CGRP and PIEZO2 in DRG. Consistent with our hypothesis, we observed an increase in both total and sensory nerve innervation in the knee joints of the M_a_ group, which may underline enhanced OA-associated pain. By integrating behavioral, histological, molecular, imaging and computational approaches, our study provides novel insights into the neurobiological mechanisms of OA pain and highlights potential sex-dependent prognostic and therapeutic strategies.

## Material and methods

### Experimental animals

A total of n = 38 mice (both male and female) were used for the study. Wild-type (WT) C57BL/6 mice were grouped as young (3-5 months) and aged (20-22 months). Ethical approval was obtained by the University Committee on Animal Resources (UCAR Protocol #2019-008). For details, see Suppl. Methods.

### Pressure application measurement (PAM) assay

Knee hyperalgesia was measured using a PAM device (Ugo Basile, Comerio, Italy) (19). Briefly, mice were scruffed and an increasing magnitude of force was applied to the hind limb knee joint at a rate of 30 g/s, up to a maximum of 450 gf. Knee joint withdrawal and/or vocalization were used as indicators of the pain bearing threshold. Results are expressed as knee withdrawal threshold (gf). The assessment was performed by an investigator blinded to the experimental groups.

### Safranin O/Fast Green (Saf-O/FG) staining

Mouse knee joints were harvested, fixed in 10% neutral buffered formalin for 72 h, and decalcified in 14% ethylenediaminetetraacetic (pH 7.4) for 2 weeks at room temperature (20). Tissues were paraffin-embedded, sectioned sagittally at 7 µm, deparaffinized, rehydrated through graded ethanol to water, and stained with Saf-O/FG to visualize articular cartilage.

### DRG isolation and immunolabeling

Lumbar dorsal root ganglia (L3–L5) were collected from young and aged mice following CO₂ euthanasia, fixed in 4% paraformaldehyde, cryoprotected, embedded, and sectioned at 10 µm. Sections were immunostained with primary and secondary antibodies (Supplementary Table1 and 2) and counterstained with DAPI. Images were acquired using a Zeiss Axio Imager 2 microscope and analyzed in ImageJ. PIEZO2 expression was quantified by threshold-based integrated density measurements of individual neuronal soma, normalized to the M_y_ group. The proportion of CGRP-immunoreactive neurons was calculated as a percentage of total PGP9.5⁺ neurons, with full methodological details provided in the **Supplementary Methods**.

### iDISCO+ volume immunostaining of mouse knee joints

Whole-mount volume immunostaining and optical clearing of mouse knee joints were performed using an iDISCO+-based protocol with minor modifications (16). Following fixation and decalcification, samples were processed for methanol dehydration, autofluorescence bleaching, immunolabeling with primary and secondary antibodies, and organic solvent–based clearing. Cleared knee joints were rendered optically transparent in dibenzyl ether prior to imaging (**Fig. 1B**). For methodological details, see **Supplementary Methods** and **Table 1 and 2**.

**Fig. 1.**
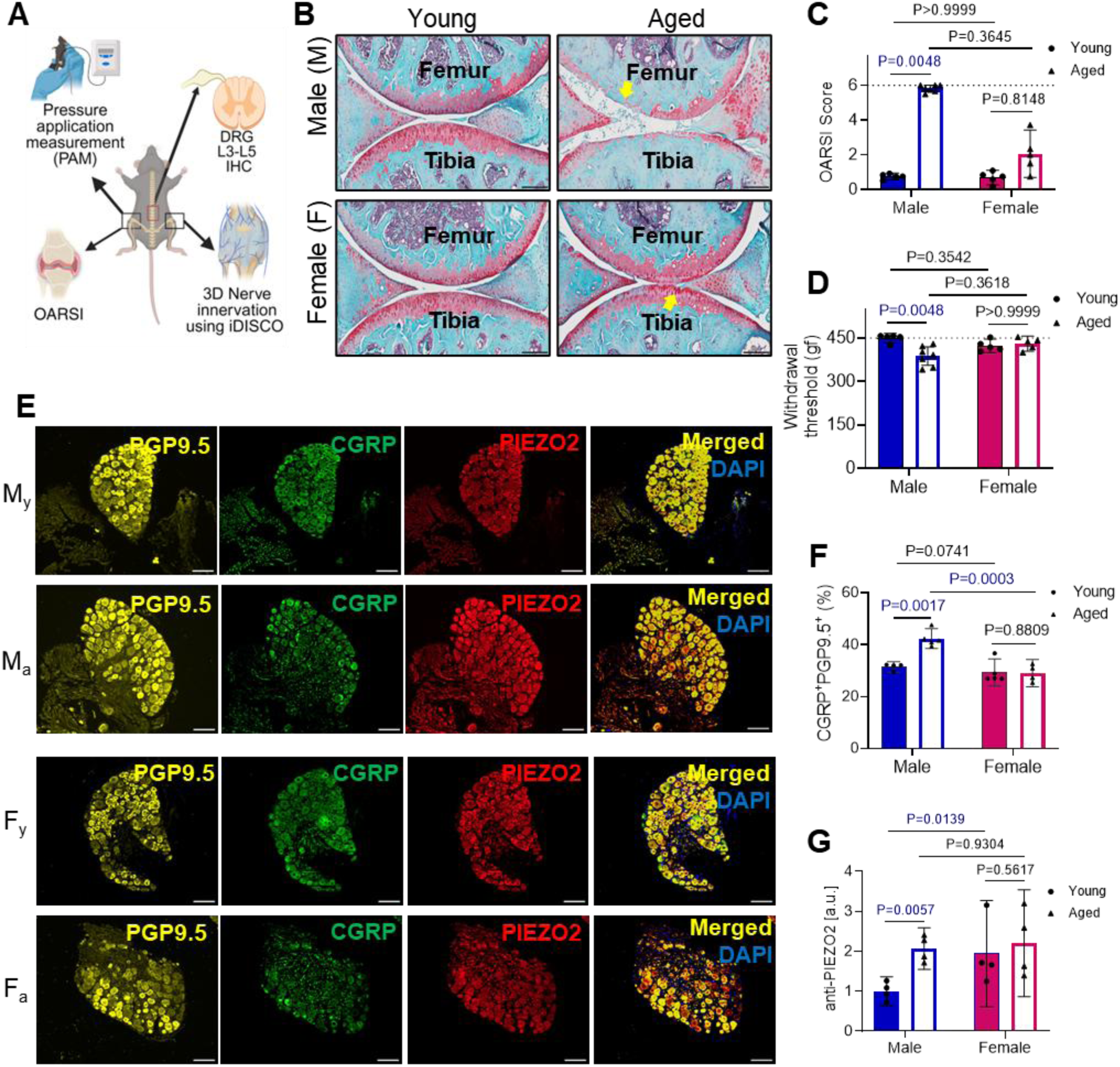
Male mice experience more severe age-associated OA relative to female mice. Age associated changes in osteoarthritis severity, pain sensitivity and expression of CGRP and PIEZO2 in the DRGs. (A) Schematic of experimental design. (B) Representative sagittal sections stained with Safranin-O/Fast-Green. Blue arrows point degenerated articular cartilage. Scale bars, 200 µm. (C) OARSI scores of the articular cartilage (n = 5-7 mice/group). (D) Quantitative analysis of knee hyperalgesia using the Pressure application measurement (PAM) (n = 5-7 mice/group). (E) Representative immunofluorescent images of PGP9.5, CGRP and PIEZO2 staining in L3-L5 DRGs. Scale bar, 100 μm. (F) CGRP^+^ neuron counts (n = 4-5 mice/group) and (G) anti-Piezo2 relative staining intensity (n = 4-5 mice/group) in DRG sections. Mean ± 95% CI. OARSI and PAM data were analyzed using a non-parametric Kruskal–Wallis test followed by Dunn’s multiple-comparison post hoc test. CGRP and Piezo2 data were analyzed using two-way ANOVA with sex and age as factors followed by uncorrected Fisher’s LSD post hoc tests.

### Light sheet microscopy of iDISCO-cleared mouse knees

Mouse knees stained and cleared via the iDISCO method described above were imaged on a Luxendo Multi-View Selective-Plane Illumination Microscope (MuVi-SPIM, Bruker) in the single detection, 4X octagon. Samples were positioned in a 3D printed holder for imaging in the frontal plane. Dual-sided illumination was performed with a series of diode laser lines (488 nm, 561nm, 642 nm) and a pair of 4X illumination objectives (Nikon, 0.2 NA). The detection arm consisted of a 4X objective (Olympus, 0.28 NA), a 1X zoom lens, corresponding bandpass filters (BP500-530 nm, BP580-627 nm, BP 655-704 nm), and a sCMOS camera (Hamamatsu Orca Flash 4.0 v3). Z-stack images (pixel size of 1.46 µm/pixel) were obtained at 3 µm increments (4.4X effective magnification), and tiled image acquisition was controlled by Lux Control software (Luxendo, v.4.4.4) set to ≥20% overlap (2400 µm) lateral step size.

### 3D image reconstruction and segmentation

Volumetric image reconstruction and 3D nerve segmentation were performed using Lux Processor and Imaris software. Tiled images were registered on the PGP9.5 (561 nm) channel, converted to .ims format, and rendered in three dimensions within a standardized region of interest encompassing the medial–lateral knee joint and proximal tibial epiphysis, including periarticular tissues. CGRP signal was quantified within reconstructed PGP9.5⁺ filaments and setting a fixed threshold to measure out the CGRP fibers from the overall nerve filaments. Filament metrics like length and branch points were exported for quantitative analysis. For details, see **Suppl. Methods**.

### Computational geometries of reconstructed nerve networks and data normalization

Spatial coordinates from reconstructed PGP9.5⁺, CGRP⁺/PGP9.5⁺, and TH⁺ nerve networks were exported from Imaris and analyzed using custom MATLAB scripts. 3D α-shapes were generated to define the spatial boundaries of each nerve network, and network volumes were estimated after excluding spatial outliers using proximity-based metrics and multivariate analysis. Network volume was then used to normalize total nerve length, while branch point counts were normalized to segment length. Medial–lateral nerve distribution was assessed by k-means clustering of segment coordinates, with full computational and statistical details provided in the **Suppl Methods.**

### Statistical analysis

All statistical analyses were performed using GraphPad Prism v10 (GraphPad Software, Boston, Massachusetts, USA). Biological replicate numbers (n) are indicated in the figure legends. Data normality was assessed for each dataset using the Kolmogorov Smirnov normality test, and data were log-transformed when the assumptions of normality were not met prior to statistical analysis. OARSI scores, PAM measurements, and imaging-derived x–y–z coordinate lengths were analyzed using Kruskal–Wallis tests followed by Dunn’s multiple-comparisons post hoc tests. All other outcome measures were analyzed using two-way ANOVA to assess the effects of age (young vs. aged), sex (male vs. female), and their interaction, with model assumptions evaluated by residual diagnostics. Post hoc comparisons were performed using Fisher’s LSD test. No statistical analyses were conducted for datasets with n ≤ 3. Medial–lateral nerve distribution was assessed using repeated-measures two-way ANOVA with compartment (medial vs. lateral) matched within animals, followed by Fisher’s LSD post hoc testing. For details, see **Supplementary Table 3 and 4**.

## Results

### Aging enhances knee osteoarthritis severity and nociceptive signaling in male mice

The experimental design (**Fig. 1A**) included young male (M_y_), aged male (M_a_), young female (F_y_), and aged female (F_a_) mice. Structural joint changes were assessed and OARSI scoring **Fig. 1B–C**). The M_y_ group displayed intact cartilage, whereas the M_a_ group exhibited marked loss of SafO staining, indicating proteoglycan depletion and OA progression. OARSI scores differed significantly among groups (H = 16.56, p = 0.0009, ε² = 0.75), with the M_a_ group showing significantly higher scores than the M_y_ (adjusted p = 0.0048); no age-related differences were observed in females. Knee joint pain was evaluated using pressure application measurements (PAM) **Fig. 1D**. Withdrawal thresholds differed among groups (H = 11.59, p = 0.0089, ε² = 0.64), with M_a_ group exhibiting significantly reduced thresholds compared with M_y_ (adjusted p = 0.0048), consistent with increased mechanosensitivity. No significant effects of age or sex were observed in females.

To characterize DRG neurons associated with mechanical sensitivity and joint pain, we examined expression levels of CGRP and PIEZO2 on DRG sections isolated from lumbar segments L3-L5 of M_y_, M_a_, F_y_, and F_a_ groups (**Fig. 1E**). The M_a_ group exhibited increased numbers of CGRP^+^ neurons in DRG compared to young (**Fig. 1F**). Our result revealed significant main effects of sex (F(1,6) = 44.33, p = 0.0006) and age (F(1,8) = 5.49, p = 0.0471), as well as a significant Age × Sex interaction (F(1,6) = 12.98, p = 0.0113). The M_e_ group showed a significant increase in the CGRP^+^ neurons compared to the M_y_ (95% CI: -0.0515 to -0.1800). In contrast, females showed no age-related change (95% CI: -0.05967 to -0.06811;). Additionally, PIEZO2 expression showed significant effects of sex (F(1,6) = 6.217, p = 0.0469) but not age (F(1,6) = 5.922, p = 0.0509), with no significant sex × age interaction (**Supplementary Table 3**). The M_a_ group exhibited increased PIEZO2 expression in DRG (mean difference = −0.3203, 95% CI −0.5283 to −0.1124), whereas no age-related difference was observed in females (mean difference = −0.05698, 95% CI −0.2650 to 0.1510) (**Fig. 1G**).

### Increased PGP9.5⁺ Nerve Innervation in Knee Joints of Aged Male Mice

Following iDISCO clearing and immunolabeling (**Supplemental Fig. S1**), knee joints were imaged by 3D light sheet microscopy. An anterior knee joint region of interest (ROI) spanning the medial–lateral width and extending from the inferior patellar pole to the proximal tibial epiphysis was selected (**Fig. 2A**). This region contained a dense PGP9.5⁺ nerve network distributed across the femur, tibia, infrapatellar fat pad, and adjacent soft tissues, primarily within the superficial joint capsule and closely associated with vasculature (**Fig. 2; Supplementary Movies M1-4**). ROI dimensions were consistent across all groups, with no age-or sex-related differences (**Fig. 2B**). Age-related changes in knee joint innervation were evaluated by labeling joint-innervating nerves with the pan-neuronal marker PGP9.5 and performing 3D filament reconstruction using Imaris (v10.1.1) (**Fig. 2C**). For the filament normalization, the 3D rendered PGP9.5+ nerve network defined in Imaris was imported to MATLAB, where computational geometry was used to calculate the volume of 3D boundary regions enveloping the PGP9.5+ nerve segments (**Fig. 2D**). PGP9.5^+^ nerve length revealed a significant effect of age (F(1,8) = 36.04, p = 0.0003) and sex (F(1,8) = 6.363, p = 0.0357). A significant sex × age interaction (F(1,8) = 6.998, p = 0.0295) revealed the differential effect of aging in male and female mice. M_a_ group showed a significant increase in total PGP9.5^+^ filament length (mean difference = −0.3127, 95% CI −0.4551 to −0.1704, p = 0.0003). In contrast, no significant age-related difference was observed in females (mean difference = 0.01799, 95% CI −0.1244 to 0.1604, p = 0.7923). While PGP9.5^+^ innervation did not differ between M_y_ and F_y_ groups (mean difference = −0.007684, 95% CI −0.2115 to 0.1962, p = 0.9329), the M_a_ group exhibited significantly greater innervation compared to the F_a_ group (mean difference = 0.3230, 95% CI 0.1192 to 0.5269, p = 0.0065). Additionally, PGP9.5^+^ branch points also showed a significant effect of age (F(1,8) = 21.71, p = 0.0016) and sex (F(1,8) = 7.6, p = 0.0248) and a significant sex × age interaction (F(1,8) = 10.45, p = 0.0120) revealed that age-related changes in branching differed between males and females. M_a_ group showed increase in PGP9.5⁺ branch points (mean difference = −0.5539, 95% CI −0.7860 to −0.3218, p = 0.0001), whereas no age-related change was detected in females. Branch point density was significantly higher in M_a_ than F_a_ group (mean difference = 0.5771, 95% CI 0.2629 to 0.8914, p = 0.0029), while no sex difference was observed in young animals (**Fig. 2 F-I**). The PGP9.5^+^ nerve length and branch points with the boundary volume normalization exhibited similar trends (**Supplementary Table 3**).

**Fig. 2.**
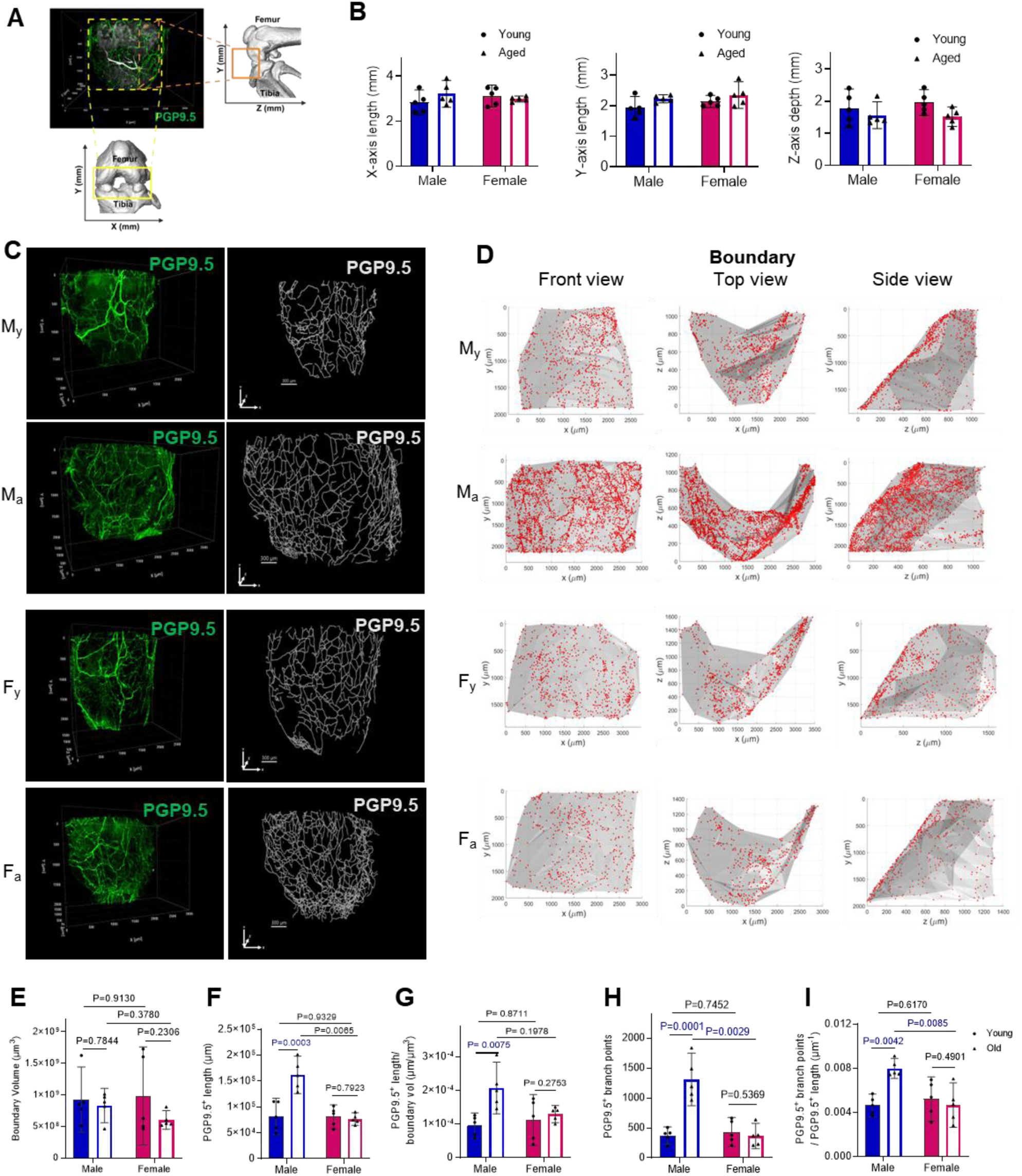
Age-associated increase in total knee joint innervation in male mice. (A) Representative three-dimensional (3D) images of knee joints labeled with anti-PGP9.5 (green) and the reconstructed µCT scans. Quantified regions are shown in yellow (x-y axis) and orange (y-z axis) boxes. (B) Quantification of the imaging coordinate length of the knee joint in young and aged male and female mice respectively in the x-axis, y-axis and z-depth. (C) Representative 3D anti-PGP9.5 (green) micrographs of the knee joint of My (young male), Ma (aged male), Fy (young female) and Fa (aged female); white line traces PGP9.5+ nerve filaments using Imaris v10.1.1 software. (D) Representative MATLAB-generated images showing the boundary volume (gray) encapsulating the PGP9.5+ segment coordinates (red) in My, Ma, Fy and Fa (E) Quantification of the total boundary volume, (F) PGP9.5+ filament length, (G) PGP9.5+ filament length/total boundary volume, (H) PGP9.5+ branch points, and (I) PGP9.5+ branch points/total filament length in young and aged male and female mice, respectively. N = 5 mice/group. Data were analyzed using two-way ANOVA to assess the effects of age and sex, followed by uncorrected Fisher’s LSD post hoc tests. Data are presented as mean ± 95%CI.

### Aged Male Mice Exhibit Increased CGRP⁺ Nociceptor Innervation in Knee Joints Compared to Young Male group

To assess age-associated changes in pain-sensory innervation in osteoarthritic knee joints, tissues were co-stained for the pan-neuronal marker PGP9.5 and the nociceptive neuropeptide CGRP (**Fig. 3A**). Given that CGRP is a secreted neuropeptide and is also present in vascular smooth muscle surrounding blood vessels, the complete nerve network was first reconstructed in 3D using PGP9.5⁺ filaments in Imaris (v10.1.1). A CGRP intensity threshold was then applied to selectively render colocalized CGRP⁺/PGP9.5⁺ sensory filaments. 3D imaging revealed dense peptidergic sensory innervation within and surrounding the knee joint, particularly in extra-synovial fat pads, including the infrapatellar fat pad, and the joint capsule (**Fig. 5A, B**). Quantitative analysis showed increased CGRP⁺/PGP9.5⁺ filament length and branching in M_a_ compared with M_y_, whereas females showed minimal age-related differences (**Fig. 3B–E**).

**Fig. 3.**
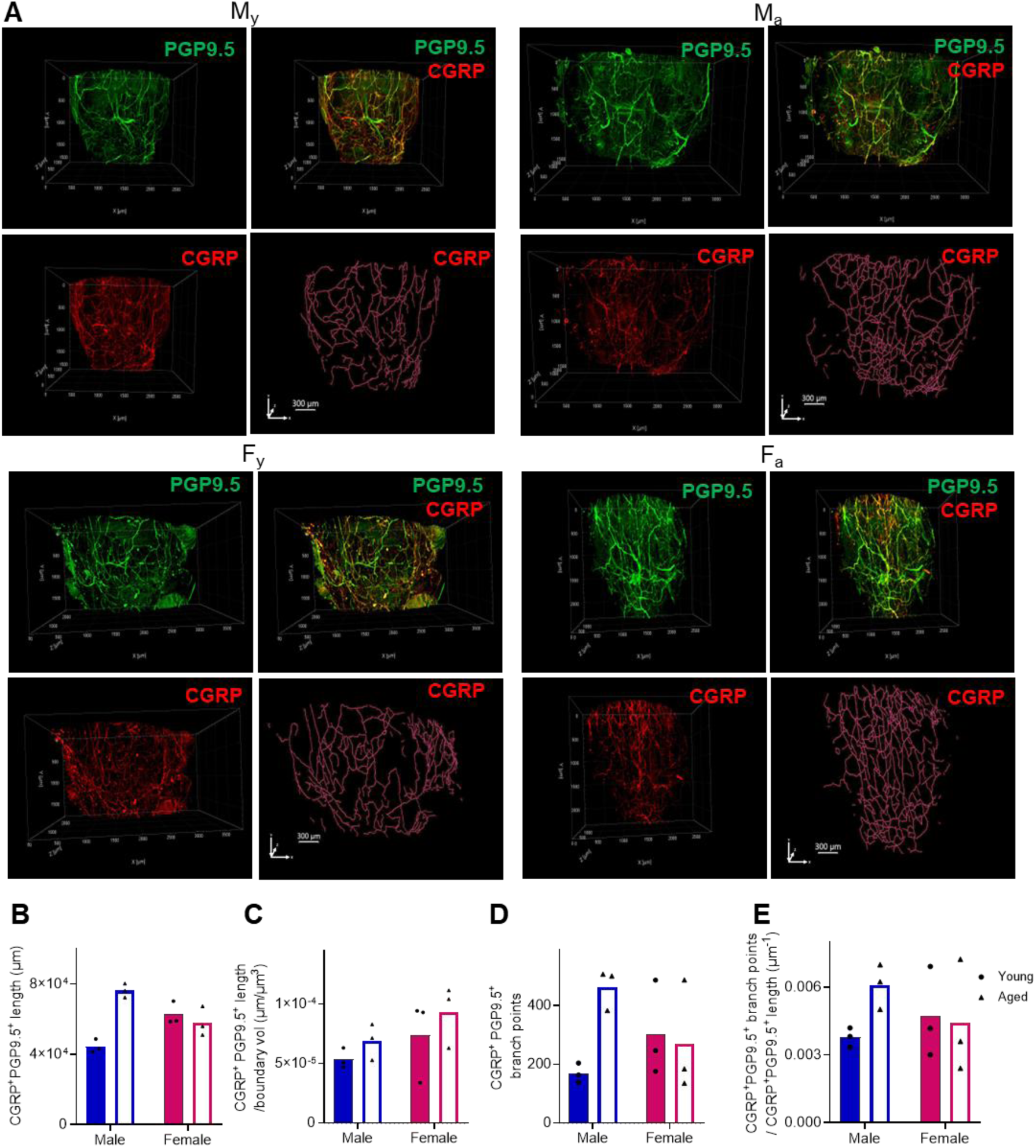
Change in CGRP⁺/PGP9.5⁺ nociceptive nerve innervation in the aging knee joints. (A) (top) Representative 3D Light-Sheet Micrographs of knee-innervating nerve fibers stained with anti-PGP9.5 (green) and anti-CGRP (red) in M_y_, M_a_, F_y_ and F_a_ ; CGRP^+^ nerve fibers are traced using IMARIS. CGRP⁺ nerve innervation was quantified by first segmenting total nerve fibers using PGP9.5 immunolabeling. A single, fixed intensity threshold was then applied uniformly across all samples to identify CGRP⁺ signal within the PGP9.5⁺ nerve segments to quantify CGRP^+^PGP9.5^+^ nerve filament. (B) Total length of CGRP^+^PGP9.5^+^ nerve filament. (C) Total CGRP^+^PGP9.5^+^ nerve filament length/boundary volume. (D) Total branch points in CGRP^+^PGP9.5^+^ nerve filaments. (E) CGRP^+^PGP9.5^+^ branch points/total CGRP^+^PGP9.5^+^ filament length. N = 3 mice/group.

**Fig. 4.**
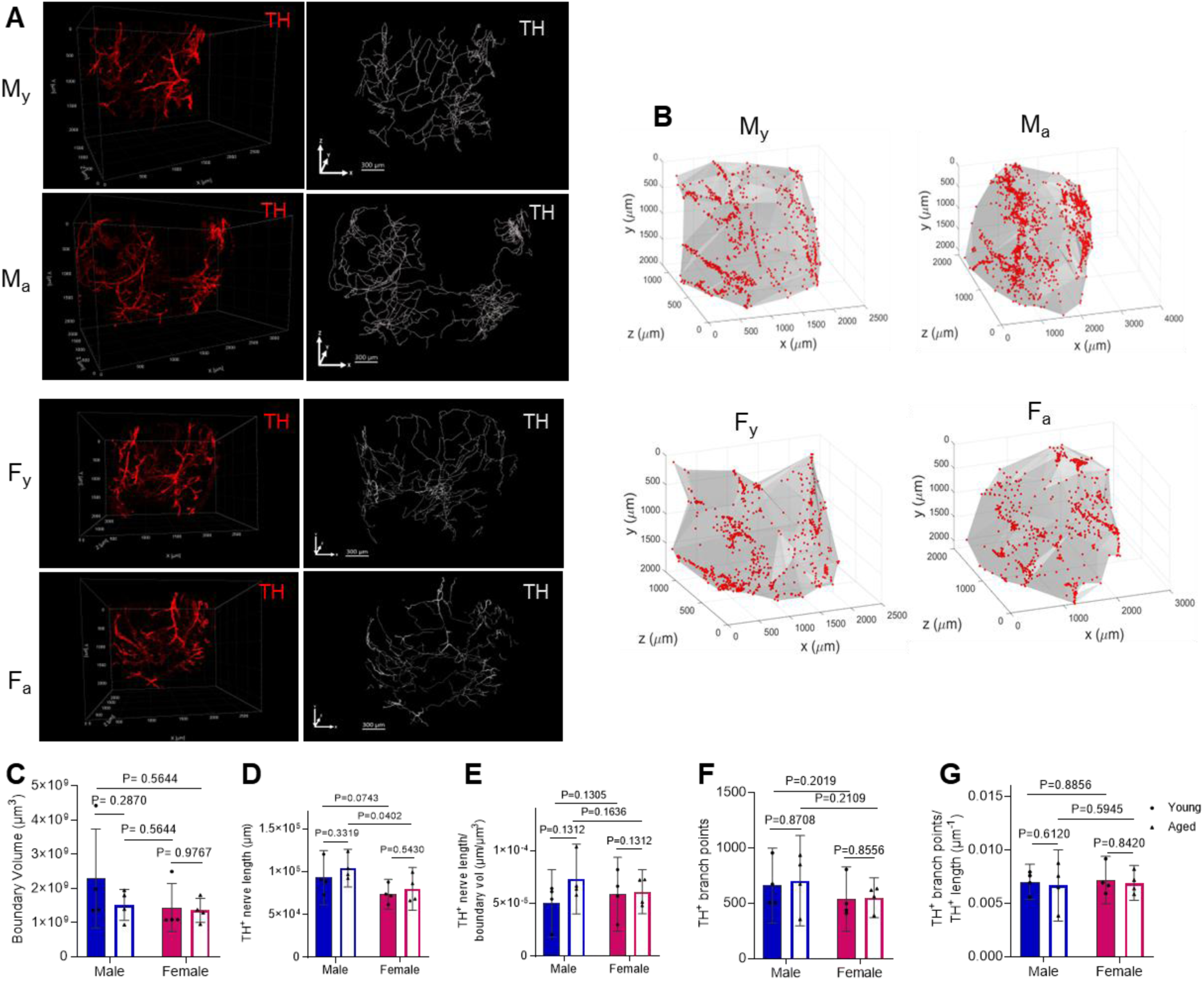
The TH⁺ sympathetic nerve network remains unchanged in male and female mice with age. (A) Representative 3D Light-Sheet micrographs with anti-TH (red) in My, Ma, Fy and Fa; TH+ nerve fibers traced using Imaris software. (B) Representative MATLAB-generated images showing the boundary volume (gray) encapsulating TH+ segment coordinates (red) in My, Ma, Fy and Fa (C) The boundary volume of TH+ filaments. (D) Total TH+ filament length. (E) TH+ filament length / boundary volume. (F) TH+ branch points (G) TH+ branch points/TH+ filament length. N = 4 mice/group. Data were analyzed using two-way ANOVA to assess the effects of age and sex, followed by uncorrected Fisher’s LSD post hoc tests. Mean ± 95%CI.

**Fig. 5.**
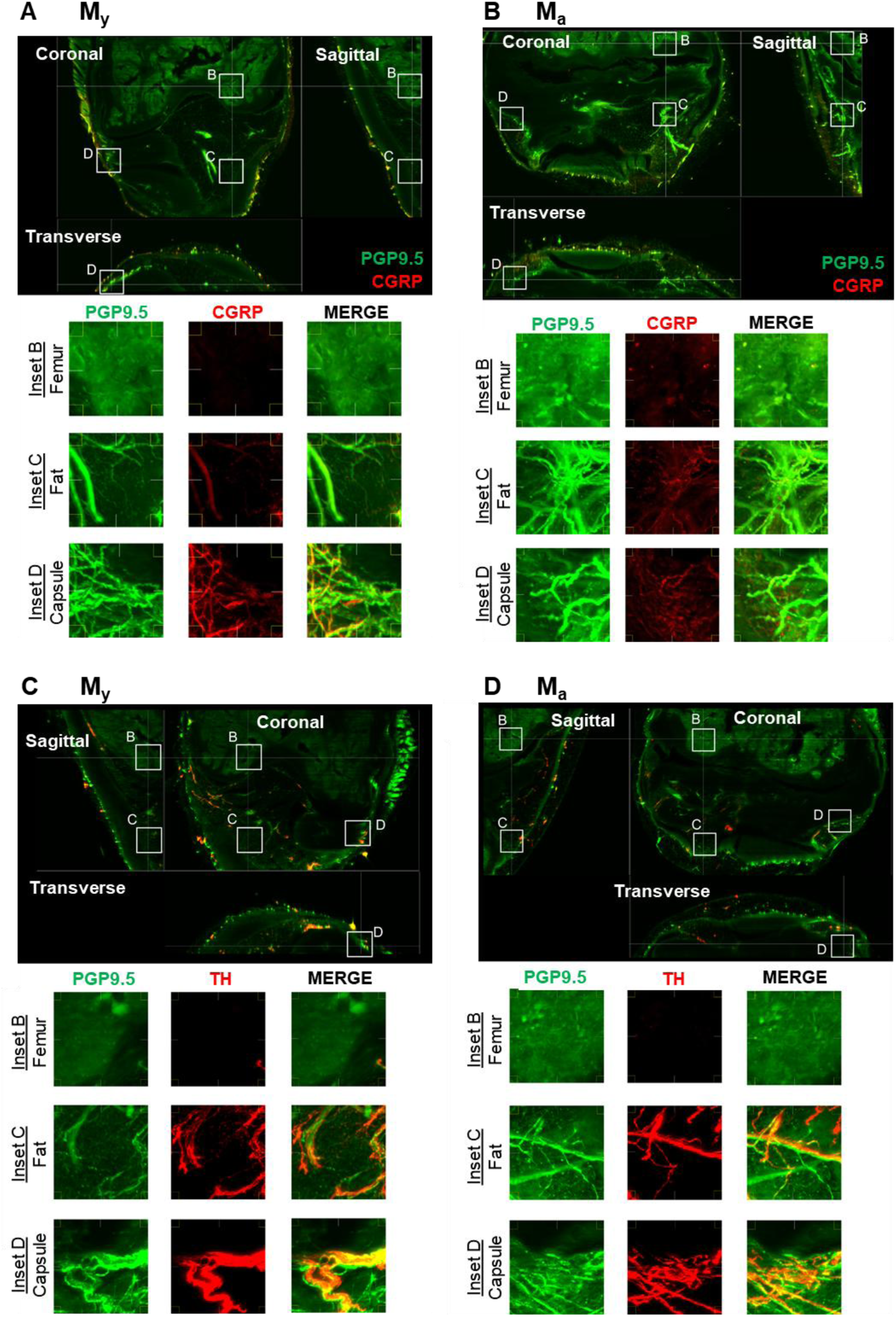
Orthogonal planes of view (coronal, sagittal and transverse) from PGP9.5/CGRP immunolabeled knee joints from (A) young male and (B) aged male mice, and from PGP9.5/TH immunolabeled knee joints from (C) young male and (D) aged male mice, depicting the depth of nerve infiltration into the various tissues of the knee joint. Insets capture the femur (inset B), infrapatellar fat pad (inset C), and joint capsule/synovium (inset D). CGRP+ sensory nerves (panels A,B) and TH+ sympathetic nerves (panels C,D) appear to innervate the extrasynovial fat and joint capsule (insets C,D), whereas minimal staining is seen in the femoral bone (inset B), which likely reflects insufficient antibody penetration into bone. White insets are 300 μm x 300 μm.

Normalization to the 3D PGP9.5⁺ network volume yielded the similar trend, confirming enhanced sensory innervation with aging in male mice.

### The Density of TH^+^ Sympathetic Nerve Fibers in Knee Joints Remains Stable During Age-Related Osteoarthritis

We next examined sympathetic nerve innervation in the anterior knee region of young and aged mice using anti-tyrosine hydroxylase (TH) labeling (**Suppl. Movies M5–8**). Representative 3D micrographs of TH⁺ sympathetic networks are shown in **Fig. 4A**. Total TH⁺ filament length and branch points were quantified in Imaris and normalized in MATLAB (**Fig. 4B**). Notably, no age-related differences were observed in TH⁺ filament length or branching in either sex (**Fig. 4C–G; Supplementary Table 3**). These findings indicate that, unlike pain-sensory nerves, sympathetic nerve density and branching remain largely unchanged with aging in osteoarthritis. In both M_y_, M_a_, and F_a_ group, TH⁺ sympathetic fibers consistently innervated extra synovial fat pads and the joint capsule (**Fig. 5C, D**).

### Spatial Distribution of Nerve Segments Between the Medial and Lateral Sides of the Knee Joint

To further characterize the spatial distribution of nerve network in knee joints, we segmented PGP9.5^+^, CGRP^+^/PGP9.5^+^, and TH^+^ filaments in medial and lateral sides using MATLAB (**Fig. 6** and **Fig. 7**). In male mice, PGP9.5⁺ nerves and CGRP⁺/PGP9.5⁺ sensory nerves were distributed across the medial and lateral sides of the knee joint without trends (**Fig. 6B, C**). However, TH^+^ sympathetic nerves were more robustly innervated in the medial aspect of the knee joint in both M_y_ and M_a_ groups (**Fig. 6D**) (**Suppl. Table 4**). In contrast, female mice revealed that PGP9.5⁺ and TH⁺ nerves were distributed across the medial and lateral sides of the knee joint without trends (**Fig. 7B, D**); yet CGRP⁺/PGP9.5⁺ sensory nerves were more robustly innervated in the lateral sides than the medial sides of the F_y_ group (**Fig. 7C**). Together, these data suggest the region-dependent joint-innervating nociceptive and sympathetic nerve remodeling over OA progression.

**Fig. 6.**
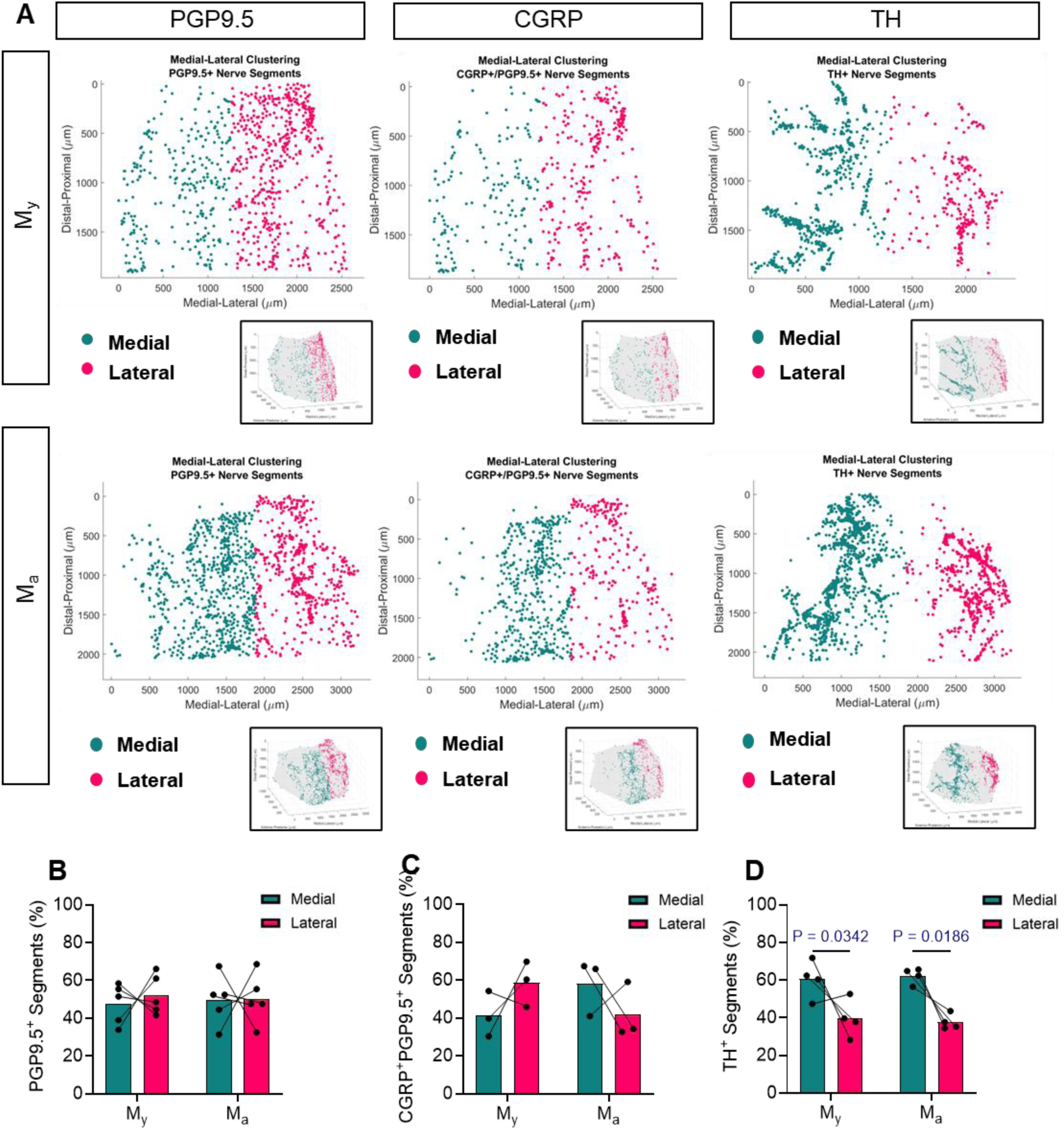
Spatial mapping of segment positions in the medial and lateral compartments of the knee joint in young and aged male mice (M_y_, M_a_). (A) k-means clustering was performed in Matlab to partition the segment position data into medial and lateral clusters for PGP9.5, CGRP and TH innervation. The percentage of segments in medial and lateral clusters was quantified for PGP9.5 (n = 5 mice/group) (B) CGRP (n = 3 mice/group) (C), TH (n = 4 mice/group) (D). Medial-lateral proportions for individual samples are connected by lines, with bars representing the mean of each compartment. Repeated measures two-way ANOVA with post-hoc Fisher’s Least Significant Difference (LSD) test was performed to statistically compare differences in medial-lateral nerve proportions across young and aged male mice, matching compartments from individual mice.

**Fig. 7.**
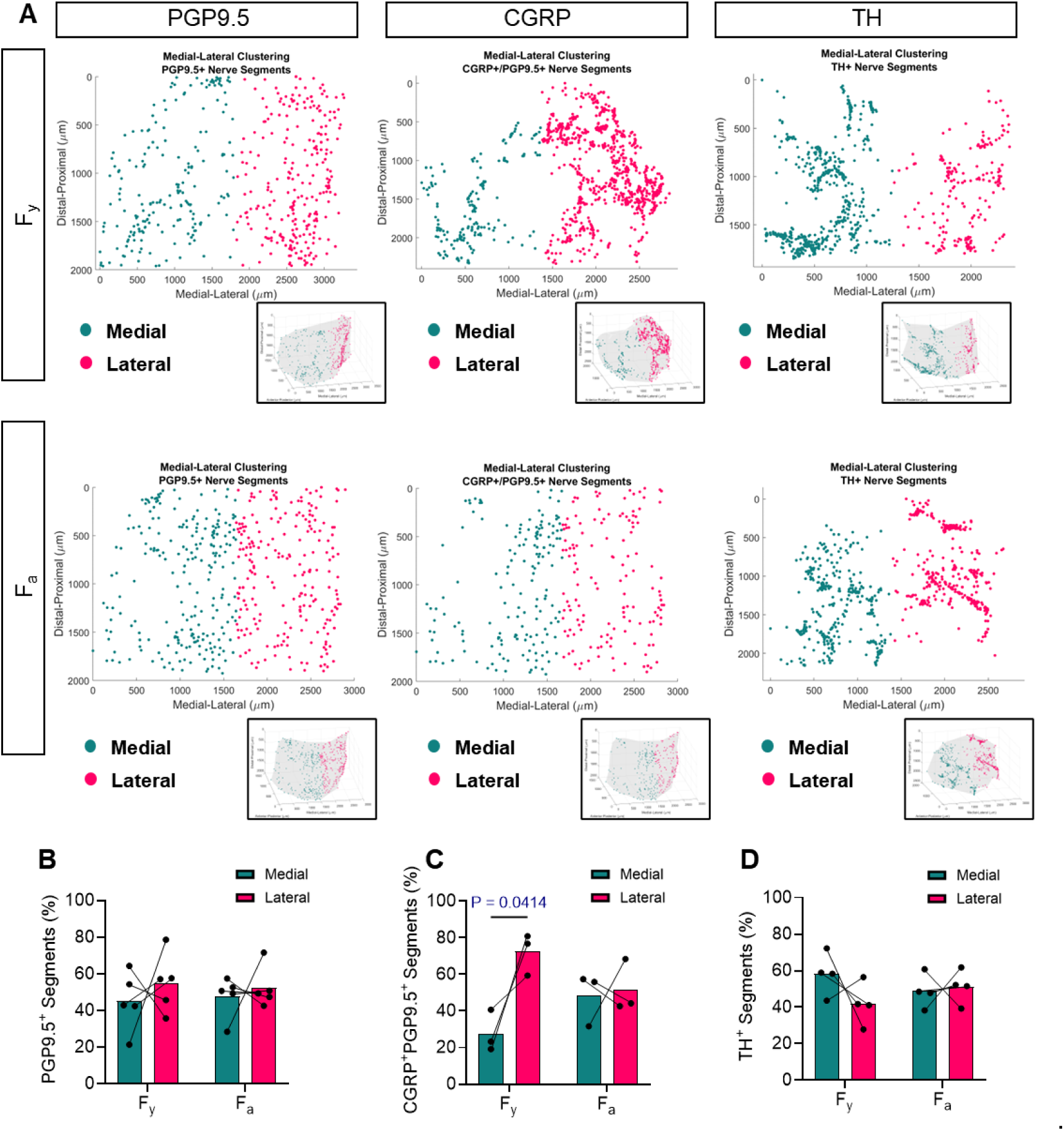
Spatial mapping of segment positions in the medial and lateral compartments of the knee joint in aged female mice with osteoarthritis compared to young female mice. (A) k-means clustering was performed in Matlab to partition the segment position data into medial and lateral clusters for PGP9.5, CGRP and TH innervation. The percentage of segments in medial and lateral clusters was quantified for PGP9.5 (n = 5 mice/group) (B) CGRP (n = 3 mice/group) (C) TH (n = 4 mice/group) (D) Medial-lateral proportions for individual samples are connected by lines, with bars representing the mean of each compartment. Repeated measures two-way ANOVA with post-hoc Fisher’s Least Significant Difference (LSD) test was performed to statistically compare differences in medial-lateral nerve proportions across young and aged female mice, matching compartments from individual mice.

## Discussion

Osteoarthritis (OA) remodels in both sensory and sympathetic nerves, contributing to nociceptor sensitization, chronic pain, and aberrant nerve sprouting (3, 21). Understanding nerve innervation patterns in aging OA is crucial, as both sensory and sympathetic fibers play pivotal roles in driving mechanical hypersensitivity within the synovial cavity (7, 14). In this study, we employed iDISCO tissue clearing combined with light-sheet microscopy to visualize 3D nerve architecture in knee joints of M_y_, M_a_, F_y_ and M_a_ mouse groups. In the examined mouse cohort, histopathological analysis showed the intact articular cartilage in knee joints of M_y_ and M_a_, while the M_a_ group exhibited severe cartilage erosion (OARSI score ∼6) and the F_a_ group with mild erosion (OARSI score ∼2.5). The PAM-based pain assessment revealed that significantly increased pain sensitivity in M_a_ than M_y_, yet not statistically significant female groups. In addition, the M_a_ group exhibited the significantly increased CGRP^+^ neurons and augmented Piezo2 in DRG compared to the M_y_ group, yet not in female groups (**Fig 1**). The increased CGRP⁺ neurons indicate enrichment of peptidergic nociceptors capable of peripheral sensitization and neurogenic inflammation, and elevated PIEZO2 expression suggests enhanced mechanical pain-associated mechanotransduction (22–25). These trends in the M_a_ group align with Geraghty et al., who reported the age-associated knee OA, mechanical pain behavior, and infiltrated immune cells in the DRG (26). However, the female groups did not exhibit enhanced pain behavior over aging in our mouse cohort. Sexual dimorphism in OA severity in aged mice may reflect variations in hormonal levels, inflammatory cytokines, and age-related reductions in spontaneous locomotor activity (27–29). Clinically, OA and associated pain are more prevalent and severe in women, who often report greater pain and functional impairment than men with similar radiographic disease (30). Conversely, murine OA models frequently display more pronounced structural damage or pain behaviors in males, with sex effects varying by model, age, and inflammatory milieu (31). These discrepancies highlight species-specific hormonal and immune differences and underscore the limitations of animal models, emphasizing cautious translation of findings from mice to humans (32).

To account for spatial variations in reconstructed nerve networks, nerve length measurements were normalized to the volume occupied by corresponding PGP9.5⁺ or TH⁺ fibers. 3D PGP9.5⁺ or TH⁺ filaments generated in Imaris were imported into MATLAB, where minimal enclosing 3D boundaries were computed (33). Quantitative analysis revealed increased total nerve length and branching in aged males, indicative of greater neural complexity, while aged females showed no significant changes. Sensory axons concentrated in the periosteum form a fishnet-like network, facilitating detection of mechanical distortions in underlying cortical bone. This likely contributes to the sharp, intense pain characteristic of OA (34, 35). Although CGRP data were analyzed descriptively due to limited sample size, patterns were consistent with PGP9.5 findings, supporting selective enrichment of nociceptive sensory fibers in aged male knee joints. Increased nerve branching and innervation may amplify the capacity to detect and transmit mechanical stimuli, potentially driving age-related mechanical hypersensitivity (18). Previous studies report divergent neural changes with aging and OA. Obeidat et al. reported reductions in CGRP⁺ nociceptors in aged OA knees, suggesting neuronal loss (14), whereas later studies documented aberrant nociceptive sprouting in synovium and subchondral bone (36). In our study, aged male mice exhibited increased innervation in the anterior knee compartment, while females showed no significant changes, highlighting sex and region-specific remodeling. Increased fiber density in males may reflect localized sprouting driven by biomechanical stress, tissue injury, or trophic/inflammatory signals, whereas females may be regulated by neuroimmune or hormonal mechanisms. Focusing on the anterior compartment rather than tissue layers underscores the importance of spatially resolved mapping when interpreting differences across studies.

Sympathetic nerves, extensively studied in the CNS, modulate sensory neurons via neurotransmitter release (37). In OA, they contribute to pain by promoting subchondral bone remodeling and influencing nociceptive signaling (38–40). Ghilardi et al. observed pronounced sympathetic innervation changes in synovium, suggesting regional susceptibility to neurovascular remodeling (41). In contrast, our anterior-knee analysis detected no age-related changes in TH⁺ fibers in either sex. Because discrete anatomical regions such as the synovium were not specifically assessed, region-specific alterations may have been missed, potentially explaining discrepancies between studies. These results indicate that sympathetic nerves in the knee remain largely stable with aging, maintaining autonomic regulation despite OA-related sensory changes.

Considering heterogeneous nerve fiber organization in aging OA (42), we examined lateral and medial compartments. Total PGP9.5⁺ fiber analysis revealed no lateral–medial differences, indicating uniform overall innervation. CGRP⁺ fibers exhibited lateral dominance in young females, absent in aged females, suggesting sensory remodeling with age. Male mice displayed no lateral–medial differences, consistent with overall increased sensory fiber density and heightened pain sensitivity. Sympathetic innervation showed medial dominance in males, potentially contributing to medial compartment OA, common in aging populations.

Knee joint innervation is spatially heterogeneous across medial, lateral, and anterior compartments, necessitating careful interpretation. Anatomical and histological studies in humans and animals reveal compartment-specific nerve input, producing unequal sensory and sympathetic fiber densities. In human osteoarthritic synovium, sensory neuropeptides such as Substance P and CGRP are denser medially, accompanied by ∼51% greater medial TH⁺ innervation (43). Obeidat et al. reported similar compartment-specific patterns in murine knees, young mice exhibited enriched nociceptive fibers in bone marrow cavities, lateral synovium, and cruciate insertions, while medial synovium had sparse innervation. Aging reduced lateral synovium and cruciate innervation, with no further loss to 52 weeks (14).

Despite the advantages of iDISCO and light-sheet microscopy for high-resolution 3D visualization, limitations exist. Bone autofluorescence can obscure signals, and maintaining consistent joint orientation is challenging, introducing variability along the z-axis. Filament segmentation was restricted to detectable regions. Our anterior joint quantifications are unlikely to compromise conclusions regarding nerve length and branching differences between ages.

Additionally, sex differences in body size and housing conditions may affect joint loading and pain independently of intrinsic OA pathology, particularly in larger aged males. We also note that the iDISCO-based workflow is adaptable to other joints and species, scaling to larger specimens requires procedure optimization. Whole-joint imaging with stitched datasets in future studies could allow comprehensive mapping.

In summary, aging induces alterations in nerve innervation and pain phenotypes in murine OA, particularly in males. These findings underscore the importance of considering sex-and region-dependent differences in OA pathophysiology. By integrating advanced imaging with computational analysis, this study provides a framework for understanding neural mechanisms driving pain in aging OA and offers a foundation for future therapeutic targeting of neural remodeling.

## Supporting information

Knee joint of Ma : A representative 3D image of the Ma (Male Aged) knee with anti-PGP9.5 and anti-CGRP.

Knee joint of My : A representative 3D image of the My (Male Young) knee with anti-PGP9.5 and anti-CGRP.

## Acknowledgments

We thank Dr. V. K. Thomas at the Center for Advanced Light Microscopy and Nanoscopy (CALMN) for help with Light Sheet Microscopy and J. Cox at the Center for Musculoskeletal Research (CMSR), Biochemistry, and Molecular Imaging Core for help with histology. The graphical figure was created using BioRender.com.

## Author Contributions

W.L. and P.K. conceptualized the study and designed the experiments. Experiments were conducted and analyzed by P.K., B.W., E.N., M,J, and Y.Z. The manuscript was written by P.K., B.W., J. J., MB, and W.L. All authors contributed to the manuscript and approved the final version.

## Role of the funding source

This research is supported by the NIH R01AR082349, NIH R35GM147054, NIH P30 AR069655, NIH T32 AR076950, NIH 1S10OD030305, and URMC Orthopedics Goldstein Award.

## Declaration of competing interests

The authors declare no competing interests.

## Notes

### Competing Interest Statement

The authors have declared no competing interest.

## References

1. Cao B, Xu Q, Shi Y, Zhao R, Li H, Zheng J, et al. Pathology of pain and its implications for therapeutic interventions. Signal Transduct Target Ther. 2024;9(1):155.

2. Malfait AM, Miller RE, Miller RJ. Basic Mechanisms of Pain in Osteoarthritis: Experimental Observations and New Perspectives. Rheum Dis Clin North Am. 2021;47(2):165–80.

3. Vincent TL. Peripheral pain mechanisms in osteoarthritis. Pain. 2020;161 Suppl 1(1):S138–S46.

4. Allen KD, Thoma LM, Golightly YM. Epidemiology of osteoarthritis. Osteoarthritis Cartilage. 2022;30(2):184–95.

5. Loeser RF. Aging processes and the development of osteoarthritis. Curr Opin Rheumatol. 2013;25(1):108–13.

6. Bartley EJ, Fillingim RB. Sex differences in pain: a brief review of clinical and experimental findings. Br J Anaesth. 2013;111(1):52–8.

7. Malfait AM, Schnitzer TJ. Towards a mechanism-based approach to pain management in osteoarthritis. Nat Rev Rheumatol. 2013;9(11):654–64.

8. Rahmati M, Nalesso G, Mobasheri A, Mozafari M. Aging and osteoarthritis: Central role of the extracellular matrix. Ageing Res Rev. 2017;40:20–30.

9. Heppelmann B. Anatomy and histology of joint innervation. J Peripher Nerv Syst. 1997;2(1):5–16.

10. Heppelmann B, Shahbazian Z, Hanesch U. Quantitative examination of calcitonin gene-related peptide immunoreactive nerve fibres in the cat knee joint capsule. Anat Embryol (Berl). 1997;195(6):525–30.

11. Jimenez-Andrade JM, Mantyh PW. Sensory and sympathetic nerve fibers undergo sprouting and neuroma formation in the painful arthritic joint of geriatric mice. Arthritis Res Ther. 2012;14(3):R101.

12. Chartier SR, Mitchell SAT, Majuta LA, Mantyh PW. The Changing Sensory and Sympathetic Innervation of the Young, Adult and Aging Mouse Femur. Neuroscience. 2018;387:178–90.

13. Kuner R, Flor H. Structural plasticity and reorganisation in chronic pain. Nat Rev Neurosci. 2017;18(2):113.

14. Obeidat AM, Miller RE, Miller RJ, Malfait AM. The nociceptive innervation of the normal and osteoarthritic mouse knee. Osteoarthritis Cartilage. 2019;27(11):1669–79.

15. Adori C, Daraio T, Kuiper R, Barde S, Horvathova L, Yoshitake T, et al. Disorganization and degeneration of liver sympathetic innervations in nonalcoholic fatty liver disease revealed by 3D imaging. Sci Adv. 2021;7(30).

16. Renier N, Wu Z, Simon DJ, Yang J, Ariel P, Tessier-Lavigne M. iDISCO: a simple, rapid method to immunolabel large tissue samples for volume imaging. Cell. 2014;159(4):896–910.

17. Ko FC, Fullam S, Lee H, Chan K, Ishihara S, Adamczyk NS, et al. Clearing-enabled light sheet microscopy as a novel method for three-dimensional mapping of the sensory innervation of the mouse knee. J Orthop Res. 2024.

18. Thai J, Fuller-Jackson JP, Ivanusic JJ. Using tissue clearing and light sheet fluorescence microscopy for the three-dimensional analysis of sensory and sympathetic nerve endings that innervate bone and dental tissue of mice. J Comp Neurol. 2024;532(1):e25582.

19. Leuchtweis J, Imhof AK, Montechiaro F, Schaible HG, Boettger MK. Validation of the digital pressure application measurement (PAM) device for detection of primary mechanical hyperalgesia in rat and mouse antigen-induced knee joint arthritis. Methods Find Exp Clin Pharmacol. 2010;32(8):575–83.

20. Kimmel D, Jee WS. A rapid plastic embedding technique for preparation of three-micron thick sections of decalcified hard tissue. Stain Technol. 1975;50(2):83–6.

21. Devor M. Pain in osteoarthritis: Driven by intrinsic rather than extrinsic joint afferents and why this should impact treatment. Interv Pain Med. 2024;3(1):100381.

22. Madar J, Tiwari N, Smith C, Sharma D, Shen S, Elmahdi A, et al. Piezo2 regulates colonic mechanical sensitivity in a sex specific manner in mice. Nat Commun. 2023;14(1):2158.

23. Murthy SE, Loud MC, Daou I, Marshall KL, Schwaller F, Kuhnemund J, et al. The mechanosensitive ion channel Piezo2 mediates sensitivity to mechanical pain in mice. Sci Transl Med. 2018;10(462).

24. Shin SM, Moehring F, Itson-Zoske B, Fan F, Stucky CL, Hogan QH, et al. Piezo2 mechanosensitive ion channel is located to sensory neurons and nonneuronal cells in rat peripheral sensory pathway: implications in pain. Pain. 2021;162(11):2750–68.

25. Wan Y, Zhou J, Li H. The Role of Mechanosensitive Piezo Channels in Chronic Pain. J Pain Res. 2024;17:4199–212.

26. Geraghty T, Obeidat AM, Ishihara S, Wood MJ, Li J, Lopes EBP, et al. Age-Associated Changes in Knee Osteoarthritis, Pain-Related Behaviors, and Dorsal Root Ganglia Immunophenotyping of Male and Female Mice. Arthritis Rheumatol. 2023;75(10):1770–80.

27. La Hausse De Lalouviere L, Morice O, Fitzgerald M. Altered sensory innervation and pain hypersensitivity in a model of young painful arthritic joints: short- and long-term effects. Inflamm Res. 2021;70(4):483–93.

28. Shoji H, Takao K, Hattori S, Miyakawa T. Age-related changes in behavior in C57BL/6J mice from young adulthood to middle age. Mol Brain. 2016;9:11.

29. Tran T, Mach J, Gemikonakli G, Wu H, Allore H, Howlett SE, et al. Male-Female Differences in the Effects of Age on Performance Measures Recorded for 23 Hours in Mice. J Gerontol A Biol Sci Med Sci. 2021;76(12):2141–6.

30. Tonelli SM, Rakel BA, Cooper NA, Angstom WL, Sluka KA. Women with knee osteoarthritis have more pain and poorer function than men, but similar physical activity prior to total knee replacement. Biol Sex Differ. 2011;2:12.

31. Contartese D, Tschon M, De Mattei M, Fini M. Sex Specific Determinants in Osteoarthritis: A Systematic Review of Preclinical Studies. Int J Mol Sci. 2020;21(10).

32. Franke M, Mancino C, Taraballi F. Reasons for the Sex Bias in Osteoarthritis Research: A Review of Preclinical Studies. Int J Mol Sci. 2023;24(12).

33. Selle M, Kircher M, Schwennen C, Visscher C, Jung K. Dimension reduction and outlier detection of 3-D shapes derived from multi-organ CT images. BMC Med Inform Decis Mak. 2024;24(1):49.

34. Castaneda-Corral G, Jimenez-Andrade JM, Bloom AP, Taylor RN, Mantyh WG, Kaczmarska MJ, et al. The majority of myelinated and unmyelinated sensory nerve fibers that innervate bone express the tropomyosin receptor kinase A. Neuroscience. 2011;178:196–207.

35. Matsuo K, Ji S, Miya A, Yoda M, Hamada Y, Tanaka T, et al. Innervation of the tibial epiphysis through the intercondylar foramen. Bone. 2019;120:297–304.

36. Obeidat AM, Ishihara S, Li J, Adamczyk NS, Lammlin L, Junginger L, et al. Intra-articular sprouting of nociceptors accompanies progressive osteoarthritis: comparative evidence in four murine models. Front Neuroanat. 2024;18:1429124.

37. Ma Z, Wan Q, Qin W, Qin W, Yan J, Zhu Y, et al. Effect of regional crosstalk between sympathetic nerves and sensory nerves on temporomandibular joint osteoarthritic pain. Int J Oral Sci. 2025;17(1):3.

38. Guan Z, Liu Y, Luo L, Jin X, Guan Z, Yang J, et al. Sympathetic innervation induces exosomal miR-125 transfer from osteoarthritic chondrocytes, disrupting subchondral bone homeostasis and aggravating cartilage damage in aging mice. J Adv Res. 2025;69:245–60.

39. Jiang W, Jin Y, Zhang S, Ding Y, Huo K, Yang J, et al. PGE2 activates EP4 in subchondral bone osteoclasts to regulate osteoarthritis. Bone Res. 2022;10(1):27.

40. Jiao K, Niu LN, Li QH, Ren GT, Zhao CM, Liu YD, et al. beta2-Adrenergic signal transduction plays a detrimental role in subchondral bone loss of temporomandibular joint in osteoarthritis. Sci Rep. 2015;5:12593.

41. Ghilardi JR, Freeman KT, Jimenez-Andrade JM, Coughlin KA, Kaczmarska MJ, Castaneda-Corral G, et al. Neuroplasticity of sensory and sympathetic nerve fibers in a mouse model of a painful arthritic joint. Arthritis Rheum. 2012;64(7):2223–32.

42. Morgan M, Thai J, Nazemian V, Song R, Ivanusic JJ. Changes to the activity and sensitivity of nerves innervating subchondral bone contribute to pain in late-stage osteoarthritis. Pain. 2022;163(2):390–402.

43. Saito T, Koshino T. Distribution of neuropeptides in synovium of the knee with osteoarthritis. Clin Orthop Relat Res. 2000(376):172–82.

